# Adaptation in plant genomes: bigger is different

**DOI:** 10.1101/196501

**Authors:** Wenbin Mei, Markus G Stetter, Daniel J Gates, Michelle C Stitzer, Jeffrey Ross-Ibarra

**Affiliations:** Dept. of Plant Sciences, University of California, Davis, CA 95616; Center for Population Biology, University of California, Davis, CA 95616; Genome Center, University of California, Davis, CA 95616

## Abstract

Here we have proposed the **functional space hypothesis**, positing that mutational target size scales with genome size, impacting the number, source, and genomic location of beneficial mutations that contribute to adaptation. Though motivated by preliminary evidence, mostly from *Arabidopsis* and maize, more data are needed before any rigorous assessment of the hypothesis can be made. If correct, the functional space hypothesis suggests that we should expect plants with large genomes to exhibit more functional mutations outside of genes, more regulatory variation, and likely less signal of strong selective sweeps reducing diversity. These differences have implications for how we study the evolution and development of plant genomes, from where we should look for signals of adaptation to what patterns we expect adaptation to leave in genetic diversity or gene expression data. While flowering plant genomes vary across more than three orders of magnitude in size, most studies of both functional and evolutionary genomics have focused on species at the extreme small edge of this scale. Our hypothesis predicts that methods and results from these small genomes may not replicate well as we begin to explore large plant genomes. Finally, while we have focused here on evidence from plant genomes, we see no *a priori* reason why similar arguments might not hold in other taxa as well.

---

Since their origin 160 million years ago, flowering plants have rapidly diversified into more than 300,000 species, adapting to a striking array of habitats and conditions. In this time, flowering plants have dominated some of the most diverse and extreme environments, harnessed a number of specialized biotic interactions in order to ensure successful pollen and seed dispersal, and adapted to meet the demands of agriculture. Given their diversity and importance, a considerable body of research has been devoted to understanding plant adaptation (Tiffin and Ross-Ibarra, 2014), but the relative importance of the various factors that may impact the process of adaptation are still not well understood. For instance, transitions in polyploidy and mating system have long been considered plausible drivers of flowering plant adaptation (Soltis et al., 2009; Goldberg and Igić, 2012), but both are also associated with evolutionary dead ends (Mayrose et al., 2011; Igic and Busch, 2013). As another example, while the effective size of plant populations is expected to correlate with estimates of the efficiency of natural selection, empirical support for this prediction is mixed (Strasburg et al., 2010; Gossmann et al., 2010). A number of aspects of adaptation have received a degree of theoretical (Hermisson and Pennings, 2017; Ralph and Coop, 2010) or empirical (Strasburg et al., 2012; Anderson et al., 2013; Ågren et al., 2013; Yeaman et al., 2016) support, but it is clear we are far from understanding all of the factors underlying the process of adaptation in flowering plants.

Here we propose that genome size may play a previously under-appreciated role in determining how plants adapt. Rather than focus on the mechanisms of genome size variation (Lynch et al., 2016; Lynch and Conery, 2003) or the adaptive significance of genome size itself (Grime and Mowforth, 1982; Bilinski et al., 2017), our **functional space hypothesis** predicts that interspecific differences in genome size may affect the process of adaptation by changing the number and location of potentially functional mutations. Below, we outline an argument for why existing differences in genome size — spanning more than three orders of magnitude across flowering plants (Gaut and Ross-Ibarra, 2008) — may lead to differences in “functional space”, and what this implies about adaptation in plant genomes.

## Larger genomes have more functional space

Mutation rates are typically reported at the level of individual nucleotides, but unless these rates change in larger genomes, plants with more nucleotides will inevitably be subject to more mutations *per genome* in each generation. And because variation in gene number across plants is not substantial (Bennetzen et al., 2005), most of the additional mutational input in larger genomes occurs outside of coding sequence.

But not only nucleotides mutate. By definition, large genomes are a consequence of insertion of additional base pairs. In plants, diploid genome size expansion is often the result of amplification of transposable elements (TEs). In addition to the nucleotides contributed by their insertion, the presence of TEs can generate variation in gene expression and regulation and even generate genome rearrangements (Chuong et al., 2016; Lisch, 2013). Beyond TEs, structural variation such as inversions are likely to be abundant in large genomes due to their association with unstable intergenic DNA. Inversions have often been found to play an oversized role in plant adaptation, from perenniality in monkeyflower (Lowry and Willis, 2010) to flowering time in maize (Navarro et al., 2017) and *Boechera* (Lee et al., 2017) to fecundity under drought in *Arabidopsis* (Fransz et al., 2016). Other structural variation is likely more common in larger genomes as well. Gene movement is frequent in large genomes such as wheat and barley (Wicker et al., 2011), frequently due to the action of transposable elements (Morgante et al., 2005). Variation in gene copy number is also common (Żmieńko et al., 2014), and has been identified as the source of a number of different adaptations (Pham et al., 2017; Prunier et al., 2017).

Together, nucleotide changes and structural changes combine to contribute an increased influx of new variation in large plant genomes. Because some portion of these mutations will have phenotypic consequences, this new variation provides greater mutational “space” on which selection can act. While clearly most mutational changes are unlikely to impact function, especially in intergenic regions, if even a small fraction of this mutational input in larger genomes impacts phenotype, selection can act to maintain or remove these mutations in populations. Indeed, the quanitity of expressed intergenic sequence appears to scale with genome size (Lloyd et al., 2017), and machine learning approaches predict a substantial minority of these sequences are likely functional (Lloyd et al., 2017). And though evidence of functionality is difficult to come by for most species, substantially more loci associated with phenotypic variation are found in intergenic regions far from genes (Figure 1) in the large genome of maize than the small *Arabidopsis* genome.

We envision that new mutations in intergenic regions are most likely to impact function if they affect regulatory sequence. Regulatory sequences across the genome can be identified by their signature of open chromatin via nuclease (MNase-seq or DNase-seq) or transposase (ATAC-seq) accessibility (e.g. Oka et al., 2017). Evidence that open chromatin may be a useful proxy for functional sequence can be found in maize, where open chromatin makes up less than 1% of the genome but nucleotide variation in such regions explains more than 40% of phenotypic variation across traits (Rodgers-Melnick et al., 2016). For the additional mutational input in larger genomes to be functional, we expect to find more of the intergenic sequence in large genomes as open chromatin and ”functional space”. In line with these expectations, Maher et al. (2017) show that total accessible chromatin space increases as genome size increases across four species ranging almost an order of magnitude in genome size (p=0.11, adjusted *R*^2^ = 0.11; Figure 2a), and that the largest increase in chromatin between species is in intergenic regions. Although chromatin accessibility data prepared in different labs and from different tissues makes direct comparison difficult, aggregated data from a broad range of species is also suggestive of increases in non-exonic open chromatin as genome size increases (Figure 2b).

**Figure 1:**
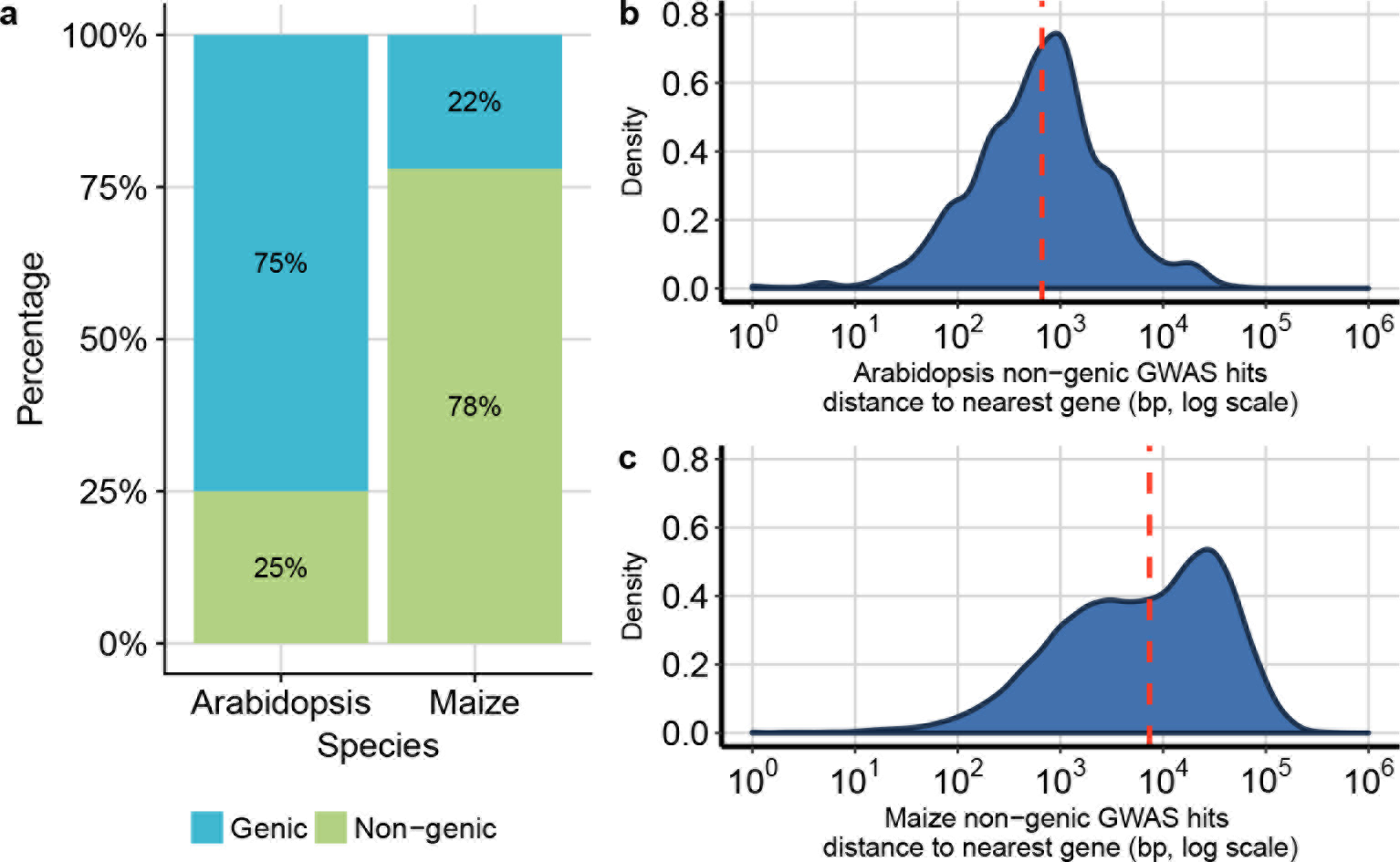
Genomic distribution of loci identified in genome-wide association studies in *Arabidopsis* and maize. *Arabidopsis* GWAS hits are top associations from the AraGWAS catalog (https://aragwas.1001genomes.org), and maize GWAS hits are the curated results of the nested association mapping population from maize diversity project (https://www.panzea.org) (Wallace et al., 2014). (**a**) Proportion of significant GWAS hits in genic (exon and intron) and non-genic regions. (**b-c**) Density plots of non-genic GWAS hits in (**b**) *Arabidopsis* and (**c**) maize. Dotted red lines indicate medians.

In total, larger genomes have more mutational input, most of which occurs in intergenic regions. Some proportion of this mutational input leads to additional open chromatin, increasing the “functional space” of large genomes.

## Large genomes adapt primarily through mutations in regulatory regions

One common approach to study adaptation polarizes synonymous and nonsynonymous substitutions using sequence comparison to an outgroup. Because nonsynonymous mutations are more likely to be functional, an observed excess of nonsynonymous substitions compared to expectations from synonymous sites likely reflects the effects of positive selection.

**Figure 2:**
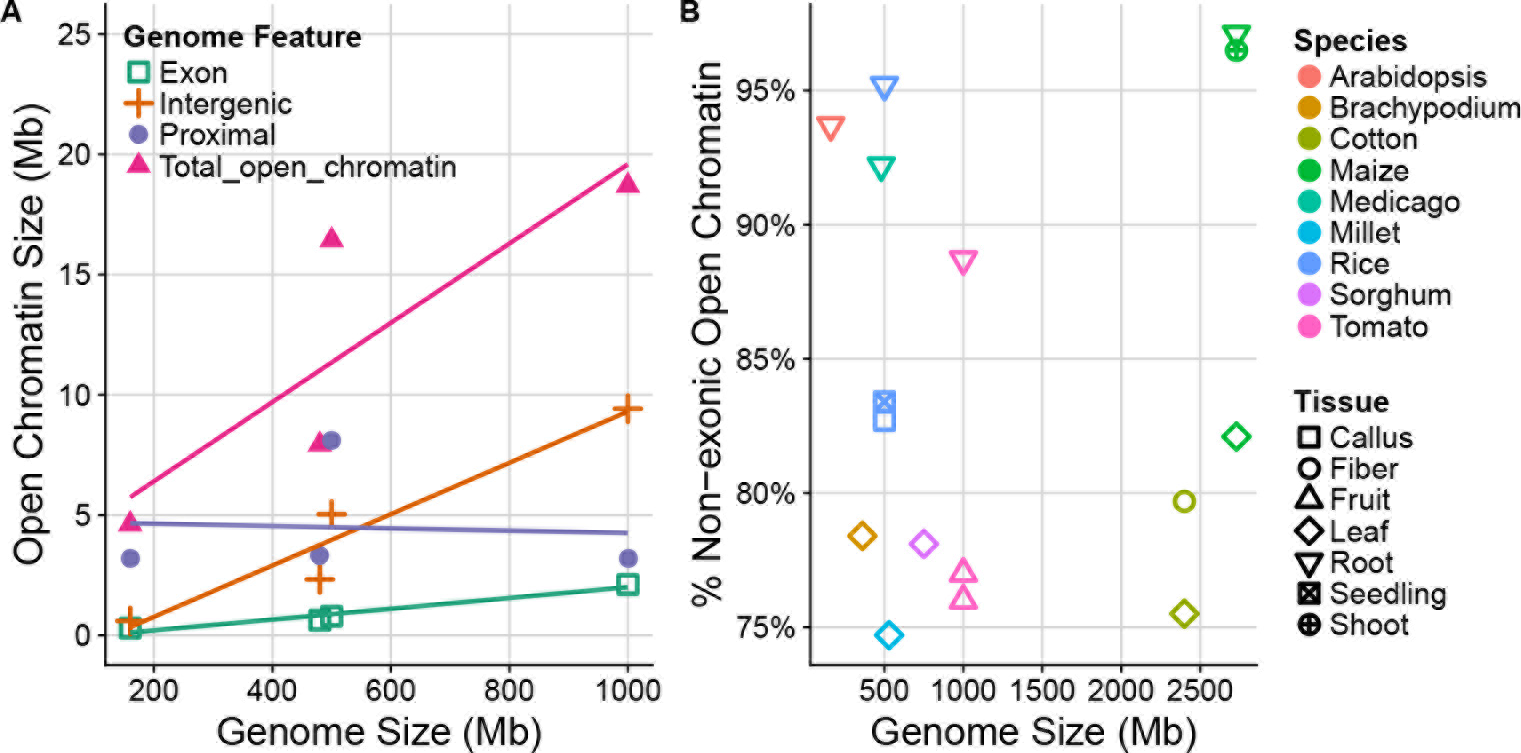
Open chromatin and genome size. (**a**) The total amount of open chromatin scales with genome size in four species *Arabidopsis thaliana, Medicago truncatula, Solanum lycopersicum* and *Oryza sativa* (Maher et al., 2017). Intergenic regions are defined >2kb from the transcription start site and >1kb from the transcription termination site. (**b**) Percentage of open chromatin outside of exons scales with genome size across many species. References and details of analyses for this figure can be found at https://github.com/RILAB/AJB_MutationalTargetSize_GenomeSize

The proportion of adaptive nonsynonymous substitutions (α) is used to infer the strength or efficiency of natural selection (Eyre-Walker, 2006). But as most of the additional functional mutations in large genomes occur in regulatory regions outside of genes (Figure 2), as genomes increase in size, a larger proportion of adaptive variation should occur outside genes and enrichment for nonsynonymous substitutions should decrease. Limited evidence for this effect can be found in two studies of environmental adaptation, where putatively selected loci are enriched for nonsynonymous mutations in the small *Arabidopsis* genome (Hancock et al., 2011) but for noncoding sequence near genes in the larger teosinte genome (Pyhäjärvi et al., 2013). We thus predict that, all else being equal, plants with larger genomes should exhibit lower *α* values than species with smaller genomes. After correcting for phylogeny, we indeed observe such a negative slope (p=0.12; Figure 3), and a negative correlation is also evident within the only two well-sampled genera available (*Helianthus* and *Pinus*). The effect we predict may be relatively weak overall, however, as a number of other factors such as effective population size and population structure (Gossmann et al., 2010) have also been suggested to impact α. Published data of α for species with small genome size are biased towards species with a small effective population size — and thus less efficient natural selection — which might influence the correlation between α and genome size in our meta-analysis (Figure 3).

## Sweeps are softer in larger genomes

At the onset of a new selective pressure, adaptation can either proceed using genetic variation currently segregating in the population or by acting upon *de novo* beneficial mutations as they enter the population. If mutations are limited, adaptation occurs predominantly by new beneficial mutations, which rapidly reach high frequency and leave a footprint of reduced local genetic diversity known as a “hard sweep” (Smith and Haigh, 1974). A number of other modes of adaptation are possible, however, and often collectively referred to as “soft sweeps”. If substantial functional variation currently exists in the population, adaptation can make use of existing variants and affect surrounding genetic diversity to a lesser degree (Hermisson and Pennings, 2017). Similarly, if the influx of new beneficial mutations across multiple loci is sufficiently high, adaptation can occur via partial sweeps that increase the frequency — but not fix — multiple independent beneficial mutations (Hermisson and Pennings, 2017).

**Figure 3:**
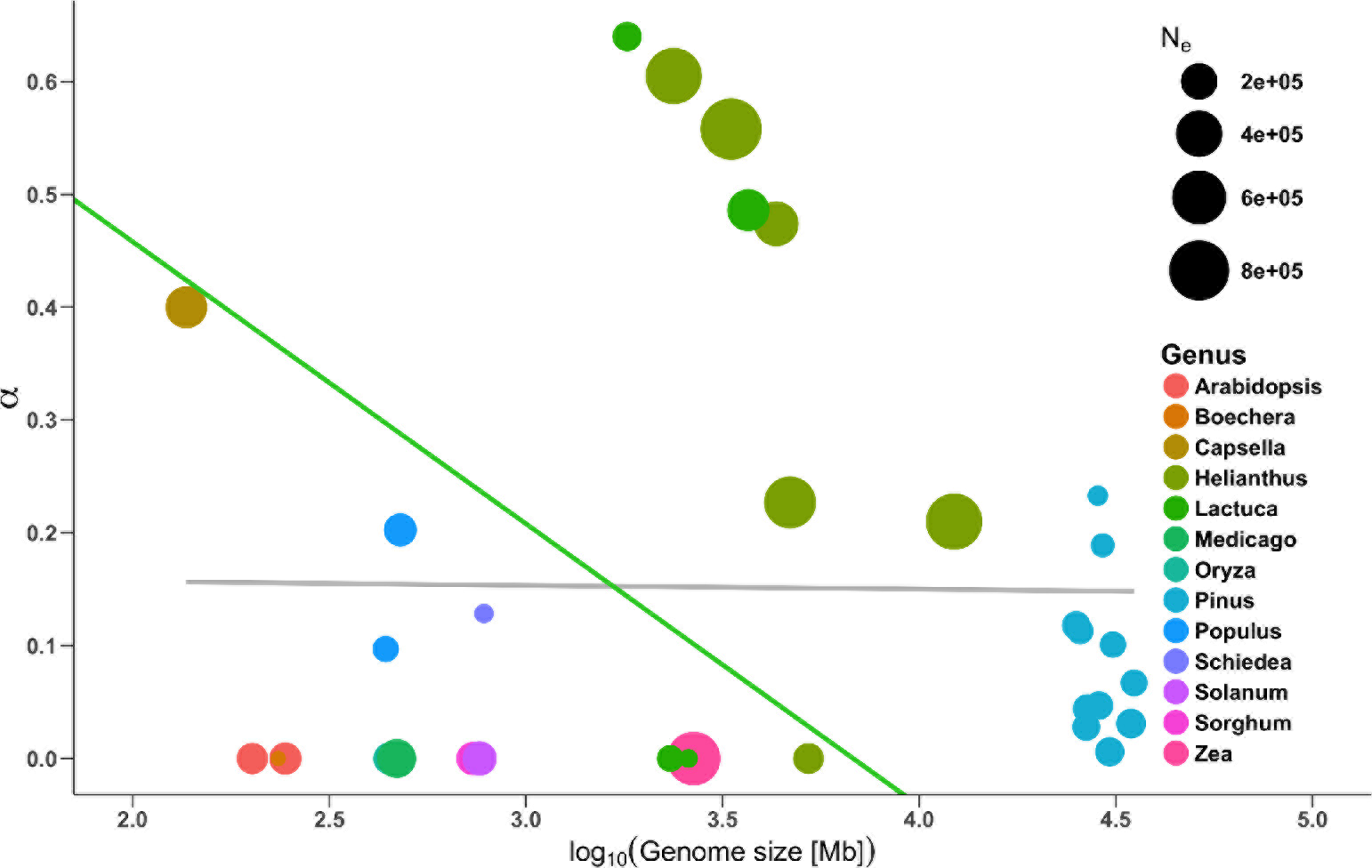
Pattern of adaptive substitutions and genome size. The proportion of adaptive substitutions (*α*) across plants with different genome sizes. Colors represent genera and point sizes show estimates of the effective population size (N_*e*_). Lines represent regression analyses performed with (green) and without (black) phylogenetic correction. References and details of analyses for this figure can be found at https://github.com/RILAB/AJB_MutationalTargetSize_GenomeSize

A key component in determining which of these processes take place as adaptation progresses is the mutational target size — what we term here more generally as “functional space”. More formally, the likelihood of sweeps being hard or soft depends on the population mutation rate *θ*_*b*_, which is the product of twice the effective population size (*N*_*e*_) and the genome-wide beneficial mutation rate (*U*_b_) (Hermisson and Pennings, 2017). While there is little evidence that the effective size of plant populations scales with genome size (Whitney et al., 2010), as we argue above the genome-wide rate of beneficial mutations should increase as genomes increase in size. Theory suggests that soft sweeps should predominate whenever *θ*_*b*_ > 1 (Hermisson and Pennings, 2017), leading to the prediction that adaptation from extant variation or multiple independent mutations should be more common in larger genomes. Although empirical examinations of the amount of soft and hard sweeps across populations is still scarce, patterns of diversity around nonsyonymous substitutions in the small *Capsella grandiflora* genome suggests a predominance of hard sweeps (Williamson et al., 2014), while similar analyses in the larger maize genome find little evidence for hard sweeps around nonsynonymous substitutions (Beissinger et al., 2016).

## Acknowledgments

The authors wish to acknowledge funding support from the NSF (IOS-1238014) and the USDA (Hatch project CA-D-PLS-2066-H) as well as Roger Deal, Pär K Ingvarsson, Toni I. Gossmann, and Aurélien Tellier for access to raw data. We thank Graham Coop and Stephen Wright for helpful discussion and Peter Tiffin, Vince Buffalo, the Prost journal club, and numerous other colleagues for comments on an early draft of the paper.

